# Genome-Wide Association Study Finds Multiple Loci Associated with Intraocular Pressure in HS Rats

**DOI:** 10.1101/2022.08.16.503865

**Authors:** Samuel Fowler, Tengfei Wang, Daniel Munro, Aman Kumar, Apurva S. Chitre, TJ Hollingsworth, Angel Garcia Martinez, Celine L. St. Pierre, Hannah Bimschleger, Jianjun Gao, Riyan Cheng, Pejman Mohammadi, Hao Chen, Abraham A. Palmer, Oksana Polesskaya, Monica M. Jablonski

## Abstract

Elevated intraocular pressure (IOP) is influenced by environmental and genetic factors. Increased IOP is a major risk factor for most types of glaucoma, including primary open angle glaucoma (POAG). Investigating the genetic basis of IOP may lead to a better understanding of the molecular mechanisms of POAG. The goal of this study was to identify genetic loci involved in regulating IOP using outbred heterogeneous stock (HS) rats. HS rats are a multigenerational outbred population derived from eight inbred strains that have been fully sequenced. This population is ideal for genome-wide association studies (GWASs) owing to the accumulated recombinations among well-defined haplotypes, the relatively high allele frequencies, the accessibility to a large collection of tissue samples, and the large allelic effect size compared to human studies. Both male and female HS rats (N=1,812) were used in the study. Genotyping-by-sequencing was used to obtain ~3.5 million single nucleotide polymorphisms (SNP) from each individual. SNP heritability for IOP in HS rats was 0.32, which agrees with other studies. We performed a GWAS for the IOP phenotype using a linear mixed model and used permutation to determine a genome-wide significance threshold. We identified three genome-wide significant loci for IOP on chromosomes 1, 5, and 16. Next, we sequenced the mRNA of 51 whole eye samples to find cis-eQTLs to aid in identification of candidate genes. We report 5 candidate genes within those loci: *Tyr*, *Ctsc*, *Plekhf2*, *Ndufaf6 and Angpt2*. *Tyr*, *Ndufaf6* and *Angpt2* genes have been previously implicated by human GWAS of IOP-related conditions. *Ctsc* and *Plekhf2* genes represent novel findings that may provide new insight into the molecular basis of IOP. This study highlights the efficacy of HS rats for investigating the genetics of elevated IOP and identifying potential candidate genes for future functional testing.

**Contribution to the field statement:** Glaucoma is the leading cause of irreversible blindness worldwide. Intraocular pressure (IOP) is the only known modifiable risk factor. This study describes results of the genome-wide association study (GWAS) performed in outbred rats that identifies known and novel genes involved in IOP regulation. To our knowledge, this is the first GWAS performed for IOP in a rat model. Identifying novel candidate genes in the rat model provides insight into the risk factors for glaucoma in humans and potential pharmacological targets for regulating IOP. The rat model is advantageous for studying natural variations in IOP, controlling environmental exposures, and providing easier access to tissue that can be used in phenotyping and gene expression in future studies.

## Introduction

Intraocular pressure (IOP) is a complex trait that is controlled by genetic and environmental factors. IOP equilibrium is achieved by the balance between production and drainage of aqueous humor. The ciliary epithelium lining the processes of the ciliary body secretes aqueous humor that enters the posterior chamber. Two pathways at the anterior chamber angle mediate the passive flow of aqueous humor from the eye. Overlying Schlemm’s canal (SC), the trabecular meshwork (TM) regulates the outflow of aqueous humor as part of the conventional route and is the primary outflow pathway (Gong et al., 1996). A fraction of aqueous outflow may travel via an ‘unconventional route’ that includes the ciliary muscle, supraciliary and suprachoroidal spaces and drains via the uveoscleral pathway (Bill, 1965). In the healthy eye, flow of aqueous humor against resistance generates an average IOP of approximately 15 mmHg (Millar, 1995). In primary open angle glaucoma (POAG), IOP becomes elevated, correlating with increased outflow resistance. Elevated IOP can cause optic nerve damage and vision loss in the majority of patients. Importantly, IOP is the only known modifiable risk factor for POAG. However, the genetic predisposition for elevated IOP remains poorly understood (Youngblood et al., 2019; Qassim et al., 2020). As a result, POAG remains the leading cause of irreversible, progressive vision loss across the globe. The number of people affected worldwide is believed to have reached 52.7 - 65 million in 2020 (Tham et al., 2014; Kapetanakis et al., 2016), which has led to an interest in identifying the specific players in this disease for the potential development of new targeted treatments.

Genome-wide association studies (GWAS) provide a strategy for discovering new genes involved in the progression of POAG by identification of genomic regions that harbor variants responsible for the variation in IOP. Many GWAS are performed in humans (MacGregor et al., 2018; Huang et al., 2019; Faro et al., 2021; Simcoe et al., 2022), however, it is difficult to perform studies in humans that further investigate how polymorphisms found in humans influence IOP. GWAS in heterogeneous stock (HS) rats provide a complementary approach that is more amenable to follow-up studies. Histologically, rats and humans share similar anatomical and developmental characteristics of the aqueous outflow pathway (Chen et al., 2016) making them an ideal model on which to study the pathophysiology of POAG.

Outbred HS rats were created by crossing eight inbred founder strains that have been fully sequenced, and thereafter maintaining them as an outbred population (Solberg Woods and Palmer, 2019); this study was performed using HS rats from the 73^rd^ to 80^th^ generations (Hansen and Spuhler, 1984; Chitre et al., 2020). The use of HS rats to study naturally occurring variations of IOP is beneficial in several ways. Numerous studies have demonstrated that phenotypic variation in humans is associated with both common and rare variants of varying effects (Cooke Bailey, J.N., Sobrin, L., Wiggs, J.L., 2020; Craig et al., 2020; Gharahkhani et al., 2021). In HS rats, all variants are common, due to the design of this population, which allows the identification of associated variants using smaller sample sizes than those required for human GWAS. Furthermore, humans are exposed to very different environmental variables that may influence IOP whereas HS rats have extremely similar environmental exposures. In addition, allelic effect sizes in model organisms are often larger than in humans and therefore can be detected with a smaller sample size (Flint and Mackay, 2009). Finally, tissue for phenotyping and gene expression studies can be easily collected from HS rats. The aim of this study was to perform a GWAS in the HS rat population to identify genomic regions that harbor variants responsible for the variation in IOP.

## Materials and Methods

### Study design

The 1,812 male and female HS rats that were used for this study were a part of a large project focused on the behavioral genetics of nicotine addiction (ratgenes.org/research-projects/rp2/; (Wang et al., 2018). We used two groups of HS rats: the older breeders that did not undergo any behavioral testing (Breeders cohort) and younger rats that underwent a battery of the behavioral tests, including brief nicotine exposure (Behavioral testing cohort). The Breeder cohort contained 733 rats with an average age of 170.1 +/−47.1 days, the Behavioral Testing cohort contained 1,079 rats with an average age of 66.4 +/−8.8 days **(Table 1 and Figure 1)**. IOP data was collected 10 days after behavioral testing was completed, allowing for any residual nicotine to be cleared. This study design allowed us to take advantage of the fact that the rats have been already genotyped and provided an important ethical advantage by using these animal subjects more judiciously.

**Table 1.** Description of two cohorts of HS rats used to measure IOP.

| Cohort | Males | Females | Total | Age, days<br>(mean ± SD) | Manipulations |
| --- | --- | --- | --- | --- | --- |
| Behavioral testing | 543 | 536 | 1,079 | 66.4 ± 8.8 | The rats were tested for multiple behaviors and were exposed to nicotine via self-administration for 12 days. IOP was measured 10 days after the final dose of nicotine. |
| Breeder | 363 | 370 | 733 | 170.1 ± 47.1 | Female breeders had one, sometimes two litters, and were never exposed to nicotine. |

**Figure 1.**
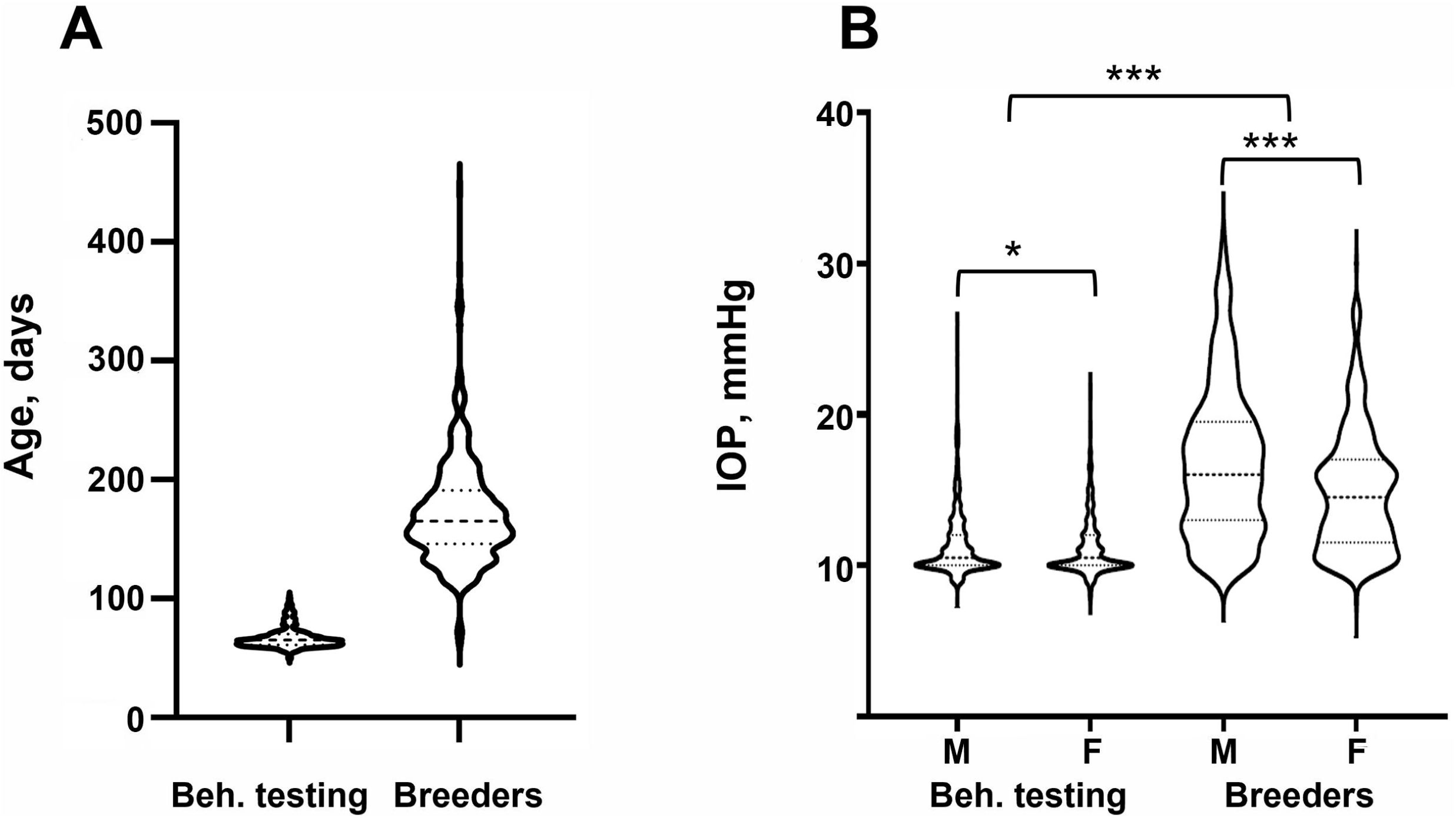
IOP in different experimental cohorts. (**A**) Age distribution for the Behavioral testing (N = 1,079) and Breeder (N = 733) cohorts. (**B**) IOP values for Behavioral testing and Breeder cohorts separated by sex. Statistical difference between sexes was tested with two-tailed *t*-test, statistical difference between Behavioral testing and Breeder cohorts was tested with Welch’s *t*-test. *denotes *p*-value <0.05, *** denotes *p*-value <0.001.

### Animals

The NIH heterogeneous stock outbred rats (HS rats) were obtained from the NMcwi:HS colony (RGD Cat# RGD_13673907, RRID:RGD_RGD_13673907). The HS rats were derived from eight inbred rat strains (ACI/N, BN/SsN, BUF/N, F344/N, M520/N, MR/N, WKY/N and WN/N) and have been maintained as an outbred population since 1984 (Hansen and Spuhler, 1984). The rats for this work were derived from generations 73 to 80. Breeders were given ad libitum access to the Teklad 5010 diet (Envigo, Madison, WI). Between 2014 to 2017, 16 batches of 50 HS rats per batch (25 males and 25 females) were sent to the University of Tennessee Health Science Center (UTHSC) at 3-6 weeks of age. These rats were then bred at UTHSC to generate offspring. Eighty-six rats were pregnant, and 26 rats were nursing during the time interval that IOP (mmHg) measurements were taken. IOP in pregnant rats (15.2 ± 4.41 mmHg) was lower than in age-matched controls (15.8 ± 5.24 mmHg). A Mann-Whitney test indicated that there was no significant difference between these groups (P = 0.5551; **Supplemental Figure 1A**). Mean IOP in nursing rats (13.9 ± 3.78 mmHg) was lower than in age-matched controls (15.5 ± 4.36 mmHg). A Mann-Whitney test indicated that there was no significant difference between these groups (P = 0.0501; **Supplemental Figure 1B**). At UTHSC, HS rats were fed the Teklad Irradiated LM-485 Mouse/Rat Diet (Envigo, Madison, WI). The rats’ offspring born at UTHSC were exposed to a battery of behavioral tests and were allowed to self-administer nicotine for 2.5 hours per day over the course of 12 days (Wang et al., 2018). A Pearson correlation was performed between IOP and total nicotine infusions received by subjects. There was a negligible correlation between the two variables and the relationship was not significant (r^2^ = 0.004, *p*-value = 0.06, N = 1079).

### IOP measurement

IOP was measured using an Icare^®^ TONOLAB tonometer (Icare, Finland). Rats were lightly anesthetized using isoflurane and then placed on a flat platform. The height of the platform was adjusted so that the center of the cornea was at the same height as the tip of the tonometer. Three consecutive measures of IOP were taken, with the maximum differences between these measures being no larger than 2 mmHg. In addition, we required the ratio of IOP from two eyes to be smaller than 1.7. Rats were given 10 minutes to rest prior to retesting if either of these requirements was not met. The phenotype used for analysis was the average of those three measurements. Both eyes were measured and the mean between the two eyes was used in the association analysis. This process was performed in the same way by the same technician for each rat to improve the consistency of readings.

### Genotyping

Genotypes for all rats were determined using genotyping-by-sequencing, as described previously (Gileta et al., 2020) The number of single nucleotide polymorphisms (SNPs) that were called before the imputation was 125,686, and the number of SNP that were called as a result of imputation and application quality filters was 3,513,494, with an estimated error rate <1%. Variants for X- and Y-chromosomes were not called for technical reasons. We removed SNPs with MAF < 0.005, a post-imputation genotyping rate < 90%, and SNPs that violated HWE with P < 1×10-10, as described in in detail previously (Chitre et al., 2020; Gileta et al., 2020).

### Heritability estimate

IOP was quantile normalized separately for males and females. Age was identified as a significant covariate for the breeders because it accounted for 3.11% of the variance. The SNP heritability (proportion of variance attributable to SNPs that were called in this population) was estimated using genome-wide complex trait analysis-genome-based restricted maximum likelihood (GCTA-GREML) software (Yang et al., 2011).

### Genome-wide association study (GWAS)

GWAS analysis was performed using Genome-Wide Efficient Mixed Model Association (GEMMA) software (Wang et al., 2016) using RGD gene dataset. Before performing GWAS, the phenotypic data was processed as follows. Each trait within a research site was quantile normalized separately for males and females. We then tested whether age explained > 2% of the variance. In Breeder cohort age explained 3.11% of variance, and was regressed out, and residuals were used for further analysis. In Behavioral testing cohort age explained < 2% of the variance. Residuals were then quantile normalized again, after which the data for each sex and cohort were pooled prior to further analysis. We did not regress out the number of nicotine infusions and pregnancy or nursing status because these groups were not significantly different, as described in “Animals” section. To account for relatedness between individuals in the HS rat population, we used a linear mixed model which can incorporate a genetic relationship matrix that models the covariance of genotypes and phenotypes among related individuals. To avoid proximal contamination, we used the “Leave One Chromosome Out” (LOCO) method (Cheng et al., 2013; Gonzales et al., 2018). To control type I error, we estimated the significance threshold by a permutation test, as described in (Cheng and Palmer, 2013). The 3.5M SNP set was LD-pruned at r^2^ < 0.99, which removed redundant SNPs to reduce computational load without impacting results. The genotype data was randomly shuffled by assigning genotype to a different individual. This random assignment scrambled the relationship between phenotype a genotype but preserved the structure within each individual genotype. GWAS was performed on the permuted data, and the maximum of the −log10 (p-values) was recorded. This step was repeated 2,500 times. The thresholds at the genome-wide significance levels 0.05 and 0.1 were respectively estimated by 95% and 90% quantiles of the 2,500 recorded maxima. The genome-wide significance threshold at the 0.1 level was −log10p = 5.67, and at the 0.05 level was −log10p = 5.95. The criteria for selection of QTLs were dependent on a SNP exceeding the significance threshold with support by at least one additional SNP within 0.5 Mb that had a p-value within [2 – log_10_(p)] units of the index SNP (Chitre et al., 2020). Regional association plots were produced using LocusZoom software (Pruim et al., 2010) using RefSeq gene dataset. The LocusZoom regional association plots show only genes from Refseq, however we also consider genes listed in the Rat Genome Database in our analysis.

### RNA-seq

Whole eyes from 53 rats (one eye per rat) were frozen immediately after each rat was killed and stored at −80°C until RNA could be extracted. During RNA extraction, eyes were first thawed to room temperature and extraocular muscles were carefully removed. RNA was extracted using the Qiagen RNeasy Midi kit following the manufacturer’s instructions. Libraries were then constructed for sequencing (150 bp paired-end reads) using the Illumina NovaSeq instrument by Novogen, Inc using the manufacturer’s protocol. This produced an average of 29.6 million read pairs per individual (standard deviation 10.1 million).

### Expression quantitative trait loci (eQTLs)

Paired-end RNA-Seq libraries for 53 rat eyes were aligned to the Rnor_6.0 genome using Spliced Transcripts Alignment to a Reference (STAR) (Dobin et al., 2013). Samples were aligned in two passes employing novel splice junctions identified in the first pass to improve alignments. The RNA-seq samples were tested for mismatches with their genotypes by inferring genotypes from RNA-seq reads and comparing them to DNA genotypes. The set of SNPs lying within exons and with MAF >= 0.2 and <10% missing values were selected from the DNA genotypes. Genotypes at these SNPs were also inferred from RNA-seq samples by counting alleles in the RNA sequences using GATK ASEReadCounter. The similarity between the genotypes from the DNA genotyping pipeline and those inferred from the RNA was computed for each rat. (McKenna et al., 2010). We compared the RNA-inferred genotype from each rat to the DNA genotype of all other rats in the same manner and found that 11 of the mismatched RNA samples were a strong match to a different DNA genotype, none of which already had a match, and could be recovered simply by relabeling the RNA samples, while the remaining two RNA samples had no high-similarity match in the DNA genotypes. Using these corrected RNA-seq/genotype pairs and removing the two remaining mismatches, we continued with 51 rat eye samples. Gene expression was quantified using RSEM, and transcripts per million values were normalized using inverse normal transformation (Li and Dewey, 2011). The first five principal components computed from the genotypes and the first 20 principal components computed from the gene expression data were used as covariates. cis-eQTL mapping was performed using tensorQTL with a cis-window size of +/-0.5 Mb (Taylor-Weiner et al., 2019). The per-gene significance thresholds were computed by tensorQTL from data permutations, and false discovery rate (FDR) testing was performed across all genes using a q-value cutoff of 0.05. We also ran tensorQTL in cis nominal mode with larger cis-window sizes for genes of interest to obtain nominal p-values for all SNPs in the plotted regions.

## Results

### 1. IOP differences in demographic groups

There was a significant difference in IOP between the Behavioral testing cohort and the Breeders cohort. Subcohort analyses revealed that males had significantly higher IOP than females in both cohorts. (**Table 2, Figure 1**). We then quantile-normalized IOP data separately for males and females within each cohort. Combining both cohorts and including both males and females allowed us to increase the power for the subsequent genetic analysis.

**Table 2.** IOP (mmHg) in different experimental cohorts.

| Cohort | Females |  |  | Males |  |  |
| --- | --- | --- | --- | --- | --- | --- |
|  | N | Avg<br>(mmHg) | std | N | Avg<br>(mmHg) | std |
| Behavioral testing | 536 | 11.2 | 2.05 | 543 | 11.5 | 2.44 |
| Breeders | 370 | 14.9 | 4.00 | 363 | 16.8 | 5.01 |

### 2. Heritability

The SNP heritability estimate for IOP was 0.317 + 0.032. The SNP heritability is expected to be somewhat lower than heritability estimated in twin studies or by using inbred strains. Heritability of IOP reported in human studies ranges from 0.39 to 0.64 (Chang et al., 2005; Carbonaro et al., 2009; Renard et al., 2010), heritability estimated using BXD mouse strains was 0.31 and 0.35 (Chintalapudi et al., 2017; King et al., 2018).

### 3. GWAS

Using the combined cohort of 1,812 rats, we performed a GWAS and identified three genomewide significant loci for IOP located on chromosomes 1, 5, and 16 (**Figure 2**). Top SNP positions and −log10(p) values are: chr1:151709252, −log10(p) = 6.000; chr5:23768259, −log10(p) = 6.742; chr16:76528844, −log10(p) = 9.221. The QQ plot is shown in Supplemental **Figure 5**. The inflation factor was 1.405. The inflation of the QQ plot (**Supplemental Figure 5**) could be due to cryptic relatedness and population stratification or true polygenic signal spread across regions of high LD. We have previously demonstrated that our methods effectively control for population structure (Gileta et al., 2022; Gonzales et al., 2018).

**Figure 2.**
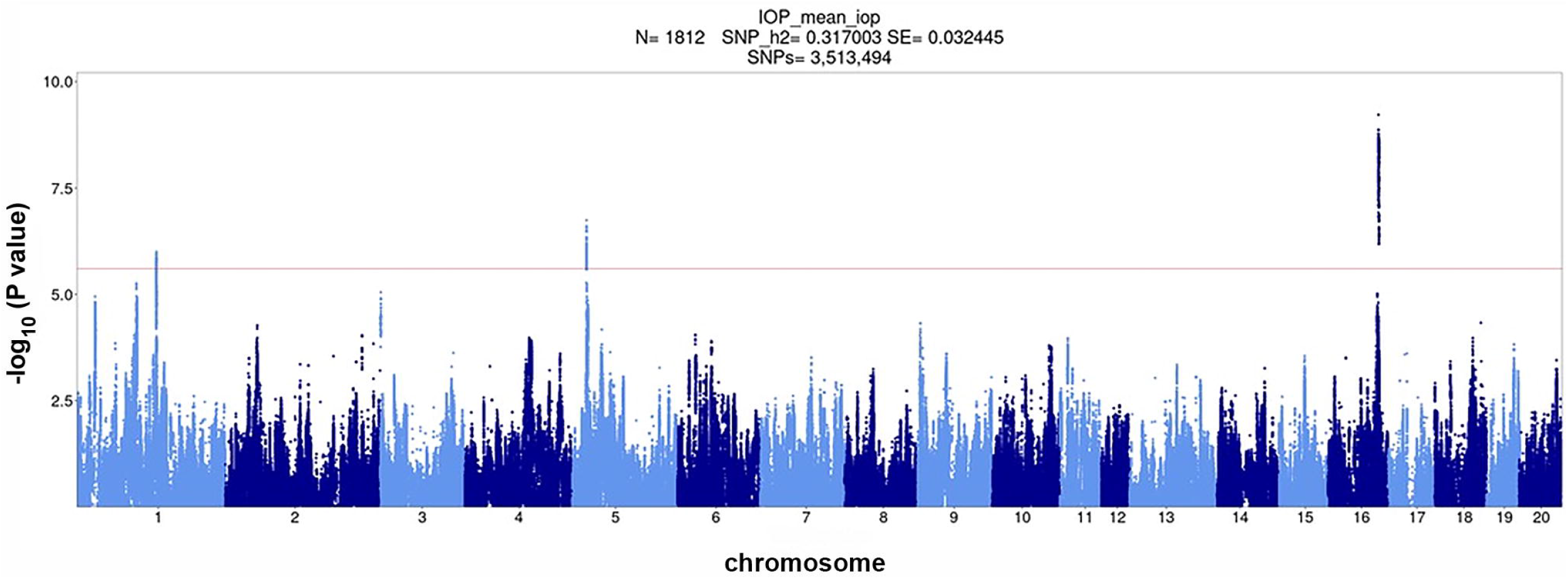
Association of ~3.5 million SNPs with IOP measured in 1,812 HS rats. The red horizontal line corresponds to genome-wide significance level 0.1 (-log10p = 5.67) the dashed red horizontal line corresponds to genome-wide significance level 0.05 (-log10p = 5.95) as determined by permutation analysis.

### 4. eQTLs

We performed eQTL analysis using 51 rat eye samples. We identified 778 cis-eQTLs that are listed in **Supplemental Table 1**. The data presented in this table can also be viewed and explored in the RatGTEx portal (ratgtex.org). Note that this portal is a resource that enables comparing gene expression in the eye with expression data for other tissues that have been produced by other researchers: for example, expression in several brain areas. **Table 3** shows expression association for the genes that are located in QTL regions identified by the GWAS analysis described in this paper. **Supplemental Figures 2, 3** and **4** show association of expression of genes, located in each QTL, with genotypes.

**Table 3.** Association of gene expression with genotypes in QTLs located on chromosomes 1, 5, and 16. These genes are located within the LD region that is defined as r^2^ > 0.6 with the top SNP for each QTL. Genes that have cis-eQTL are bolded. *denotes genes that lie just outside the interval of interest, but are included because of supporting evidence from prior publications.

| Gene name | eQTL top SNP | TSS position, bp | Log2 of Effect size | $-\log_{10}(p)$ | $-\log_{10}(p)$ threshold |
| --- | --- | --- | --- | --- | --- |
| *Tyr* | chr1:151274936 | 151,106,802 | -0.649 | 2.11 | 3.28 |
| *Grm5* | chr1:151188824 | 151,207,846 | -0.502 | 1.72 | 3.42 |
| ***Ctsc*** | chr1:151908338 | 151,918,514 | 0.117 | 3.51 | 3.37 |
| *Rab38* | chr1:152014155 | 152,072,716 | -0.158 | 1.42 | 3.45 |
| *Tmem135* | chr1:152688130 | 153,096,225 | 0.210 | 0.63 | 3.61 |
| *MGC94199* | chr5:23536473 | 23,358,464 | 6.644 | 1.25 | 3.68 |
| *LOC102555924* | chr5:24663199 | 24,230,769 | 1.005 | 0.84 | 3.46 |
| *Plekhf2* | chr5:24082110 | 24,255,922 | -0.264 | 3.38 | 3.50 |
| *Ndufaf6\** | chr5:24227315 | 24,320,804 | -0.516 | 2.38 | 3.52 |
| ***Angpt2\**** | chr16:75564376 | 75,966,480 | -0.483 | 3.62 | 3.62 |
| *Mcph1* | chr16:76216277 | 76,110,624 | 1.361 | 2.72 | 3.64 |
| *Csmd1* | chr16:78525074 | 77,152,823 | -0.112 | 0.72 | 3.48 |

### 5. Candidate gene identification

To identify candidate genes within each QTL, we considered several criteria: whether the gene was located within an interval that contains SNP in a high LD with the top SNP (r^2^ > 0.6), the presence of moderate or high impact variants located within the gene, as predicted by SnpEff (Cingolani et al., 2012), and the presence of a significant cis-eQTL detected in the eye.

Chromosome 1 contained a genome-wide significant locus with the top SNP located at position 151,709,252 bp with −log_10_(p) = 6.0 (**Figure 3**). The interval of high LD (r^2^ with the top SNP > 0.6) spanned from 151,096,891 to 152,730,623 bp. This region includes four genes: tyrosinase (*Tyr*), glutamate metabotropic receptor 5 (*Grm5*), cathepsin C (*Ctsc*), and Ras-related protein Rab 38 (*Rab38*) as well as four uncharacterized genes: *LOC102554680*, *LOC102555105*, *LOC102555017*, and *LOC365305* that are not shown on **Figure 3**. *Ctsc* is the only gene where the association between expression level and genetic variants exceeds the significance threshold (**Table 3, Supplemental Figure 2A**). The top SNP for the *Ctsc* eQTL is in strong linkage disequilibrium with the top SNP of the QTL (r^2^ = 0.985), providing supporting evidence for *Ctsc* being a candidate gene. Detailed cis-eQTL data for *Ctsc* and other genes in this region is shown in **Supplemental Figure 2A.** The *Tyr* gene is also located within the high LD region. Our eQTL data did not support *Tyr* as a candidate gene because the *Tyr* gene expression is not associated with genotypes in this region (**Table 3**). However, the *Tyr* gene contains high-impact coding and has a well-characterized variant C->T at position 151,097,606, that changes amino acid 299 from Arg to His. This modification changes the functionality of the tyrosinase protein, rendering it dysfunctional and causing the albino phenotype (Kuramoto et al., 2012). High LD (r^2^ = 0.91) between the *Tyr* C/T variant at 151,097,606 and the QTL top SNP supports the possibility that this variant is causal for the QTL on chromosome 1. In addition, the effect plot suggests that IOP tends to be higher in rats with the genotype TT than in rats with either genotype CC or CT (**Supplemental Figure 2B**), although this difference is not statistically significant. Together, these data suggest that *Tyr* is a candidate gene for IOP. *Grm5* and *Rab38* are also located in the high LD region but they do not have cis-eQTLs in this chromosomal area (**Table 3, Supplemental Figure 2A**) and they do not contain high-impact coding SNPs, therefore they were not selected as candidate genes.

**Figure 3.**
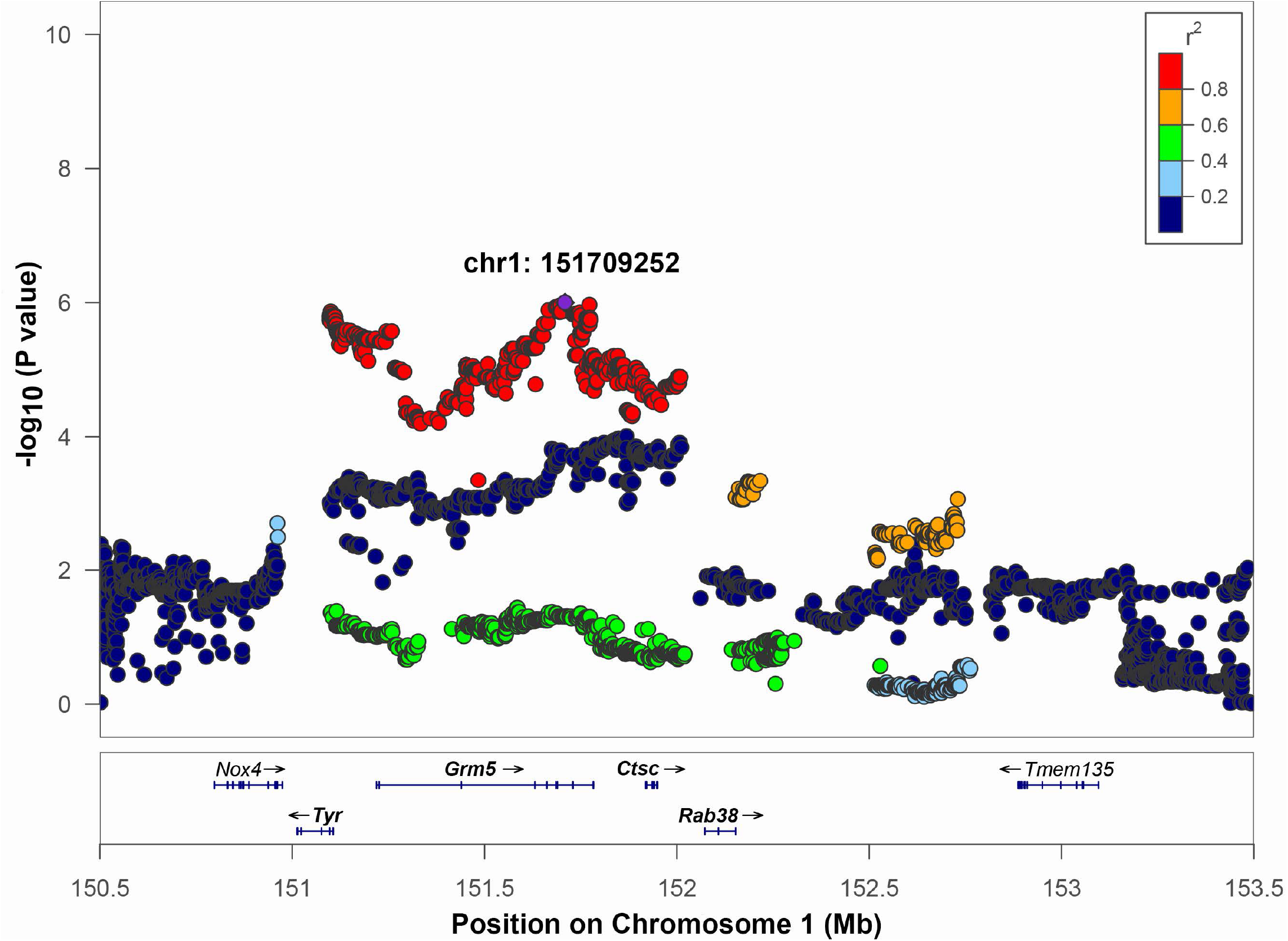
Regional association plot for QTL on chromosome 1. The x□axis shows the chromosomal position. The SNP with the lowest P value (“top SNP”) is shown in purple, and its chromosomal position is indicated on the plot. The color of the dots indicates the level of linkage disequilibrium (LD) of each SNP with the top SNP, as measured by correlation. The bottom panel indicates the genes in the region, as annotated by the NCBI Reference Sequence Database. Genes that are bolded are located in the interval of high LD of the top SNP based on a r^2^ >0.6 window.

Chromosome 5 contained a genome wide significant locus with the top SNP located at position 23,768,259 bp, −log_10_(p) = 6.742 (**Figure 4**). The LD interval of interest range was 23,426,708 to 24,262,026 bp based on a 0.6 r^2^ window. This region includes two genes: *MGC94199* and pleckstrin homology and FYVE domain containing 2 (*Plekhf2*). Also, it contains one tRNA gene: *Trnas-aga2* and four uncharacterized genes: *LOC679087, LOC100360939, LOC102555924*, and *LOC108350932* that are not shown on **Figure 4**. The regional association plot for chromosome 5 is shown on **Figure 4** with other genes labeled within the range from 23-26 Mb containing the IOP QTL. *MGC94199* does not have supporting cis-eQTL. Association analysis for *Plekhf2* expression shows that nominal *p*-value for cis-eQTL is close to the threshold (**Table 3, Supplemental Figure 3**). The gene NADH:Ubiquinone Oxidoreductase Complex Assembly Factor 6 (*Ndufaf6*) lies outside of the region of high LD with r^2^ > 0.6, but belongs to an expanded LD region with 0.4 < r^2^ < 0.6. Genes *Plekhf2* and *Ndufaf6* have been shown to play a role in conditions related to IOP, as discussed below.

**Figure 4.**
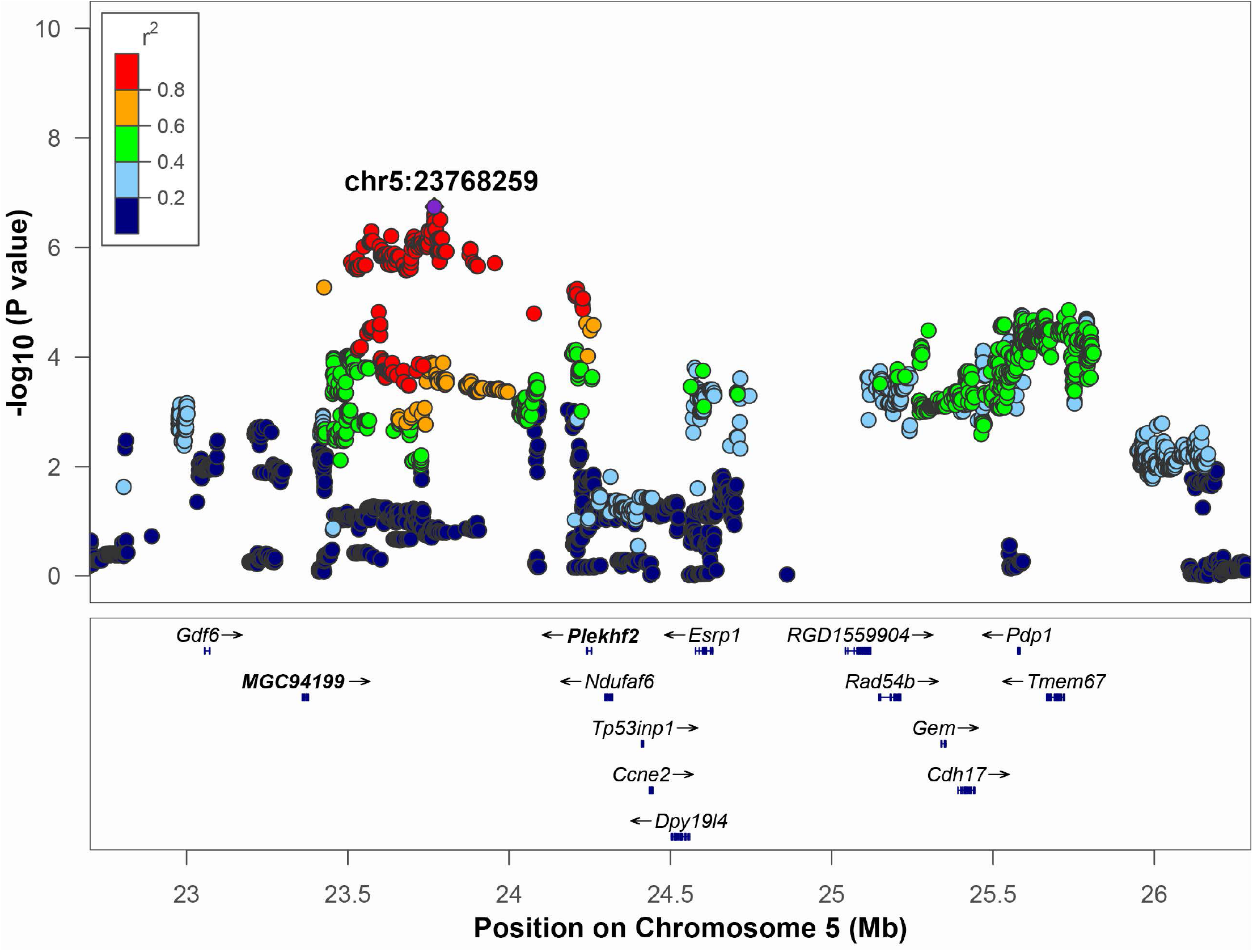
Regional association plot for QTL on chromosome 5. The x□axis shows the chromosomal position. The SNP with the lowest p-value (“top SNP”) is shown in purple, and its chromosomal position is indicated on the plot. The color of the dots indicates level of linkage disequilibrium (LD) of each SNP with the top SNP, as measured by correlation. The bottom panel indicates the genes in the region, as annotated by the NCBI Reference Sequence Database. *The *Ndufaf6* gene lies outside of the interval of high LD (r^2^ > 0.6). Genes that are bolded are located in the interval of high LD of the top SNP based on an r^2^ > 0.6 window.

**Figure 5.**
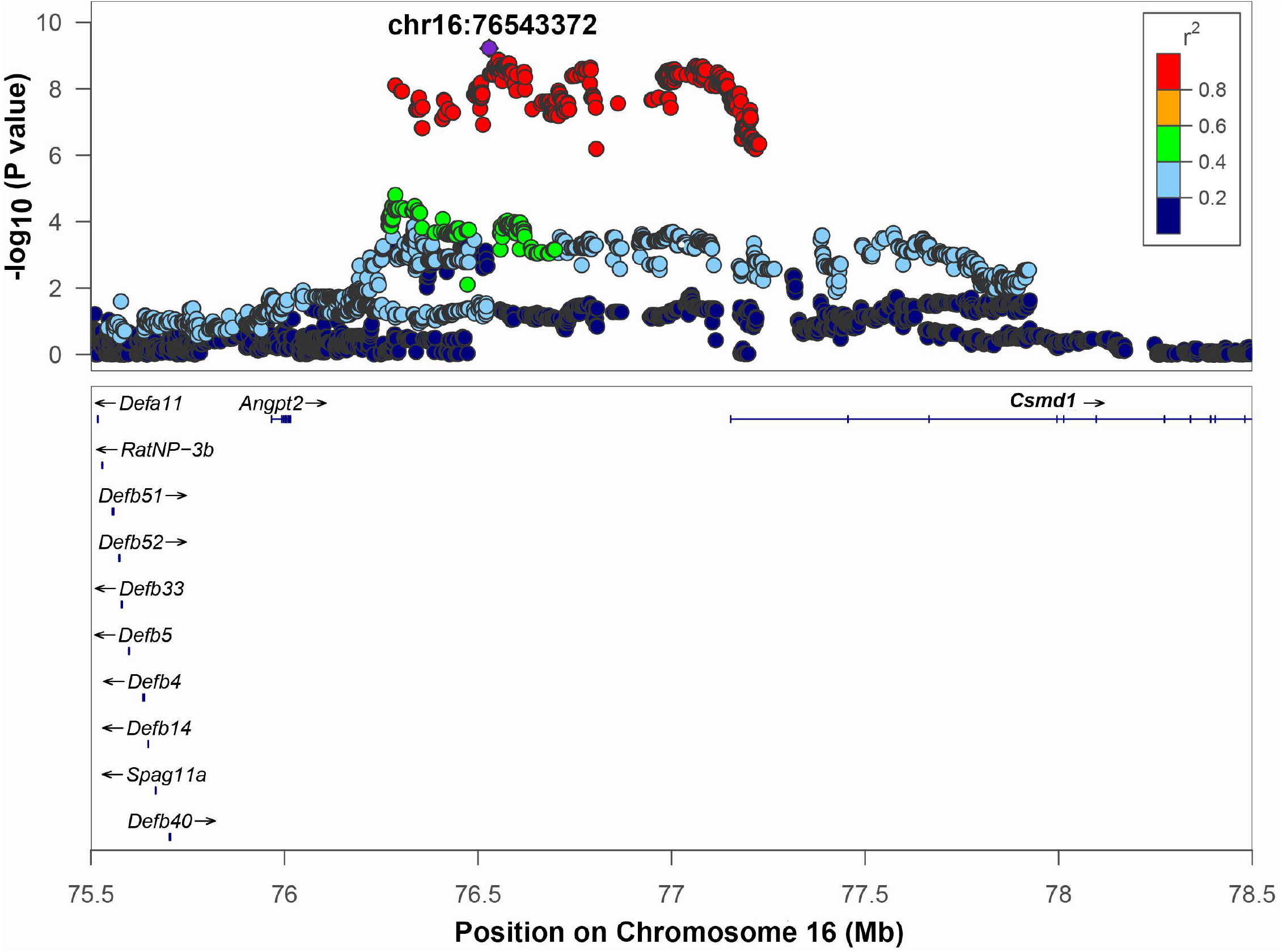
Regional association plot for QTL on chromosome 16. The x□axis shows the chromosomal position. The SNP with the lowest p-value (“top SNP”) is shown in purple, and its chromosomal position is indicated on the plot. Color of the dots indicates level of linkage disequilibrium (LD) of each SNP with the top SNP, as measured by correlation. The bottom panel indicates the genes in the region, as annotated by the NCBI Reference Sequence Database (O’Leary et al., 2016). *\*Angpt2* gene lies outside of the interval of high LD (r^2^ > 0.6). Genes that are bolded are located in the interval of high LD of the top SNP based on an r^2^ > 0.6 window.

Chromosome 16 contained a QTL with the top SNP located at position 76,528,844, −log_10_(p) = 9.221. The range of high LD was 76,286,915 to 77,225,982 bp based on a r^2^ > 0.6 window. This region includes one gene: CUB And Sushi Multiple Domains 1 (*Csmd1*) and two uncharacterized genes: *LOC108348424* and *LOC108348426* that are not shown on **Figure 5**. The regional association plot (**Figure 5)** includes a wider range 75.5 to 78 Mb. *Angpt2*, a known gene involved in IOP-related regulation (see discussion), is located close to the QTL, but is not in high LD with the top SNP. However, the p-value for the *Angpt2* eQTL is at a threshold for significance (**Table 3, Supplemental Figure 4**), suggesting that *Angpt2* expression is regulated by variants located in chromosome 16 QTL. *Csmd1* gene does not have cis-eQTL or high-impact variants, and not selected as a candidate gene.

## Discussion

GWAS is a powerful approach that allows for the identification of genome regions that harbor variants responsible for the variation in quantitative traits, such as IOP. A strength of GWAS studies is the number of subjects analyzed. This study included 1,812 HS rats, which is without precedent for genetic studies of IOP using rodents, but still relatively small compared to human GWAS. HS rats provide greater power for GWAS on a per-subject basis when compared to human GWAS due to the larger effect sizes, the larger linkage disequilibrium blocks, the absence of rare variants, and the relatively uniform environment (Solberg Woods and Palmer, 2019). These advantages have allowed us to identify multiple loci with genome-wide significance. Our study demonstrates the use of existing large-scale rodent GWAS studies to analyze additional traits that can be collected in the same population, such as IOP, in the context of a study designed to analyze behavioral and addiction traits. We identified three genome-wide significant QTLs on chromosomes 1, 5, and 16. Genes contained in these regions include those involved in IOP regulation (*Angpt2*, *Tyr*, *Ndufaf6*), and genes for which involvement in IOP regulation has not been previously described in the literature (*Ctsc*, *Plekhf2*). Our data suggest that these genes represent novel findings that may be useful for future studies of the molecular mechanisms of IOP and POAG.

GWAS for IOP have not been previously performed in rodents, but several studies have examined the genetic basis of IOP in a rodent model. (Savinova et al., 2001) demonstrated that IOP varied in 30 inbred strains, suggesting that IOP is a heritable trait in mice. A genetic study using 38 BXD recombinant inbred strains identified QTL on chromosome 8, and selected *Cdh11* as a candidate gene (King et al., 2018). In another study, IOP was measured in a panel of 65 BXD strains, identifying a QTL on chromosome 5, with *Cacna2d1* selected as a candidate gene based on systems genetic approach (Chintalapudi et al., 2017). Note that the QTL on mouse chromosome 5 identified by (Chintalapudi et al., 2017) is not synthetic with the QTL on rat chromosome 5 that was identified in our study. Research of genetic underpinning of IOP in humans include multiple GWAS studies, however, most of them have a case-control design, and are not designed to studying the genetics of the normal variation of IOP (reviewed in (Youngblood et al., 2019; Sakurada et al., 2020; Han et al., 2021)). This study is the first to use HS outbred rats in a GWAS analysis to investigate IOP.

The significant locus for IOP that is located on chromosome 1 contains four genes of interest: *Tyr*, *Grm5*, *Ctsc*, and *Rab38*. Of the positional candidates in this interval, *Ctsc* is the strongest candidate gene because it has a cis-eQTL in this region: indicating that variants in QTL region affects the expression of this gene in the eye. In the literature, *Ctsc* has not been associated directly with elevated IOP. *Ctsc* belongs to the caphesins family, which are lysosomal proteases that play a vital role in protein degradation (Yadati et al., 2020). Another cathepsin, *Ctsk*, has been shown to be involved in IOP modulation in humans via regulation of the ECM where inhibition of *Ctsk* led to decreased ECM breakdown causing decreased aqueous humor outflow and subsequent elevated IOP (Soundararajan et al., 2021). While we are not aware of any reports of *Ctsc* being associated with elevated IOP, given the functional significance of other members of the cathepsin family with IOP, this gene should be further investigated.

*Tyr* is a gene encoding tyrosinase which is involved in melanin production. *Tyr* is a positional candidate gene that is not supported by a cis-eQTL. However, *Tyr* has a missense variant C->T at position 151,097,606, that causes an amino acid change Arg299His and renders protein nonfunctional, resulting in the albino phenotype (Kuramoto et al., 2012). Our work provides novel evidence that albinism in rats may be associated with higher IOP. This is supported by the finding that albino C57BL/6J mice homozygous for a tyrosinase mutation have higher IOPs than their pigmented counterparts (Savinova et al., 2001). A synthetic human region has been identified recently in a GWAS study focused on pigment dispersion syndrome and pigmentary glaucoma (Simcoe et al., 2022). These conditions arise from abnormal dispersion of pigment from the iris pigment epithelium, where pigment accumulates in the trabecular meshwork, resulting in obstruction of aqueous humor outflow and cell atrophy. The authors identified *Grm5* and *Tyr* as positional candidate genes for this region, while *Ctsc* was outside of the QTL identified by the authors. Although we both identified *Tyr* as a potential candidate gene, our conclusion about *Grm5* and *Ctsc* differ from the Simcoe study (Simcoe et al., 2022). Notably, we evaluated healthy HS rats that did not have pathologically elevated IOP, while the human study aimed to identify variants associated with pathological conditions. It is probable that we identified different genes because different genes are involved in the regulation of normal IOP and in the regulation IOP leading to the development of pathologic conditions.

The QTL on chromosome 5 contained two genes of interest: *Plekhf2 and Ndufaf6*. Although *Ndufaf6* has a weaker LD with the top SNP (0.4 < r^2^ < 0.6), it is supported by the presence of cis-eQTL. *NDUFAF6* in humans has been associated with IOP through its role in modulating central cornea thickness, which is a known risk factor for POAG (Iglesias et al., 2018; Choquet et al., 2020). This makes *Ndufaf6* an interesting candidate for follow-up studies aimed at understanding the regulation of IOP. *Plekhf2* has not been previously associated with IOP or glaucoma. However, it is expressed in lymph node tissue and is known to regulate lymphatic flow (Fagerberg et al., 2014; Shamsara and Shamsara, 2020). These findings coupled with its physiological role makes it a plausible candidate to regulate molecular and biomechanical processes involved in the regulation of IOP, because lymphatic vessels in the ciliary body are known to participate in aqueous humor drainage from the eye (Yucel and Gupta, 2015; Yücel et al., 2018). Interestingly, *Angpt2* has also been shown to be involved in the regulation of lymphatic flow in the eye (Thomson et al., 2014), highlighting the role of the lymphatics in IOP modulation.

The QTL on chromosome 16 contains one candidate gene: *Angpt2*. *Angpt2* is located outside of the high-LD interval, however it contains a cis-eQTL, and is supported by human GWAS studies of glaucoma (MacGregor et al., 2018; Gharahkhani et al., 2021). *Angpt2* is a ligand involved in the angiopoietin-tunica interna endothelial cell kinase (ANGPT-TEK) pathway that has been shown to affect the integrity of Schlemm’s canal (Kim et al., 2017; Thomson et al., 2017).

Even with these significant findings, our study is not without limitations. First, the sample size of our population was relatively small, but was still adequate to identify three independent loci. We intend to continue measuring IOP in HS rats to increase the power of this study. Second, the limited sample size of eye tissue used for RNA expression analysis (n=51) may have reduced our ability to detect eQTLs that are causally related to the 3 loci associated with IOP. In other words, the absence of a cis-eQTL for a gene may be a false negative resulting from a small sample size. The third limitation arises from the study design. Our population was part of a large behavioral study focused on genetic influences on drug abuse-relevant behaviors. As a result, a subset of the animals was briefly exposed to moderate doses of nicotine followed by a 10-day washout period before IOP measurement. Studies are largely inconclusive regarding the effect of nicotine and smoking on IOP. It has been suggested that nicotine causes transient changes in ophthalmic artery blood flow and vasoconstriction of the episcleral veins which inhibits aqueous outflow from the trabecular meshwork causing an increase in IOP (Yoshida et al., 2014; Lee et al., 2020). However, Lee et. al determined that the increase is on average less than 1 mmHg when all additional factors were adjusted (Lee et al., 2020). Furthermore, Bahna and Bjerkedal found no association between smokers, ex-smokers, and non-smokers and no relationship to smoking habits with IOP (Bahna & Bjerkedal, 1975). Consistent with these findings, we showed that there is a negligible correlation between nicotine dosage, which is proportional to total nicotine infusions administered, and IOP in this study. Thus, the likelihood of a brief nicotine exposure causing a significant elevation of IOP after the 10-day washout period is low. The fourth limitation is related to combining animals of different ages. IOP tends to increase with age in humans, and we see a similar trend in the HS rats (**Figure 1C**). Although our study design included additional variables, such as age and nicotine exposure, the loci that we did detect are likely to be robust to the subtle differences in drug, age, and environmental differences; this approach can be viewed as a strength if the goal is to generalize our findings across populations, including humans.

In conclusion, we performed GWAS in 1,812 HS rats and identified 3 QTLs that contain 4 candidate genes. *Tyr* and *Angpt2* candidate genes have strong literature support for their role in human glaucoma-related conditions. *Ctsc* and *Plekhf2* are novel findings -- these genes have not been previously known to be involved in IOP-related molecular processes. Overall, this study is a starting point for future discoveries in the complex pathogenicity of IOP regulation that could lead to new pharmaceutical developments.

## Abbreviations

IOP: intraocular pressure
LD: linkage disequilibrium
TSS: transcription start site
HS: heterogenous stock
GWAS: genome-wide association studies
QTL: quantitative trait locus
eQTL: expression quantitative trait locus
ECM: extracellular matrix
SNP: single nucleotide polymorphism

## Acknowledgments

This work was supported by R01EY021200 (MMJ), NIDA P50DA037844 (AAP); and a Challenge Grant from Research to Prevent Blindness to the Hamilton Eye Institute.

