## Supplemental Figures for "Genome-Wide Association Study Finds Multiple Loci Associated with Intraocular Pressure in HS Rats"

### Supplemental materials.

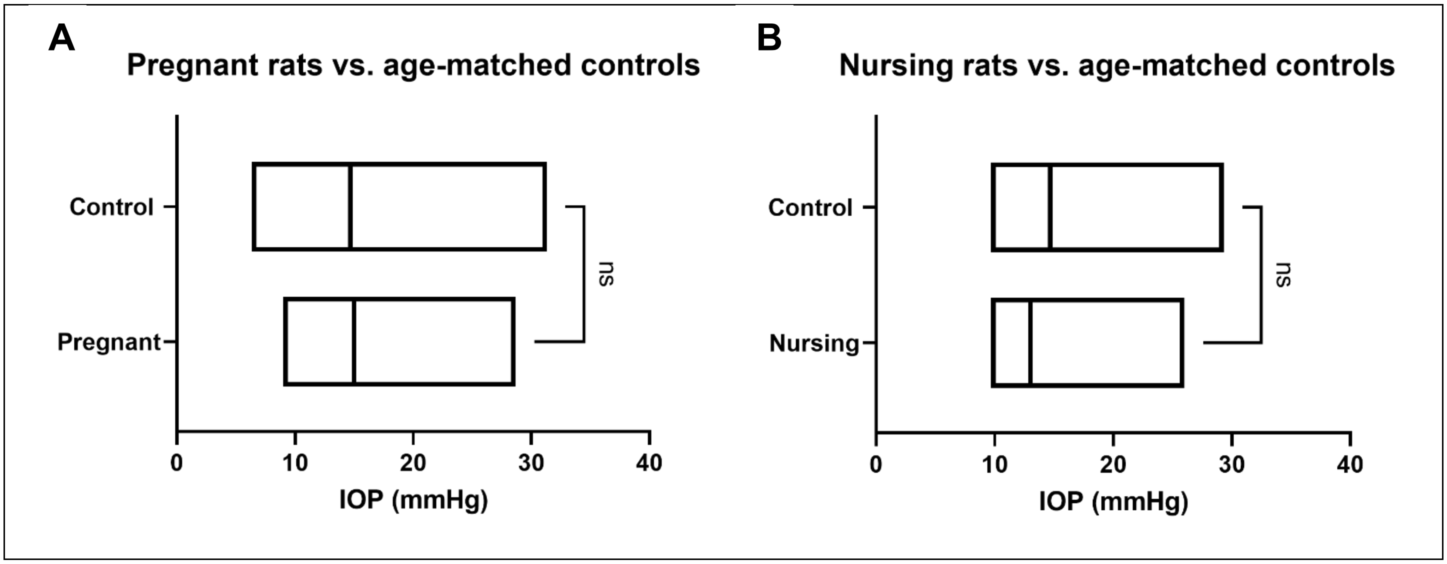


**Supplemental Figure 1.** Boxplot demonstrating IOP values from age-matched control rats vs. pregnant (n = 86) and nursing (n = 26) rats. (**A**) IOP of pregnant rats and age-matched controls. (**B**) IOP of nursing rats and age-matched controls. Number of control rats was equivalent to respective pregnant and nursing rats groups. Compared using Mann-Whitney test ($p < 0.05$). ns = non-significant.  Line shows the mean.

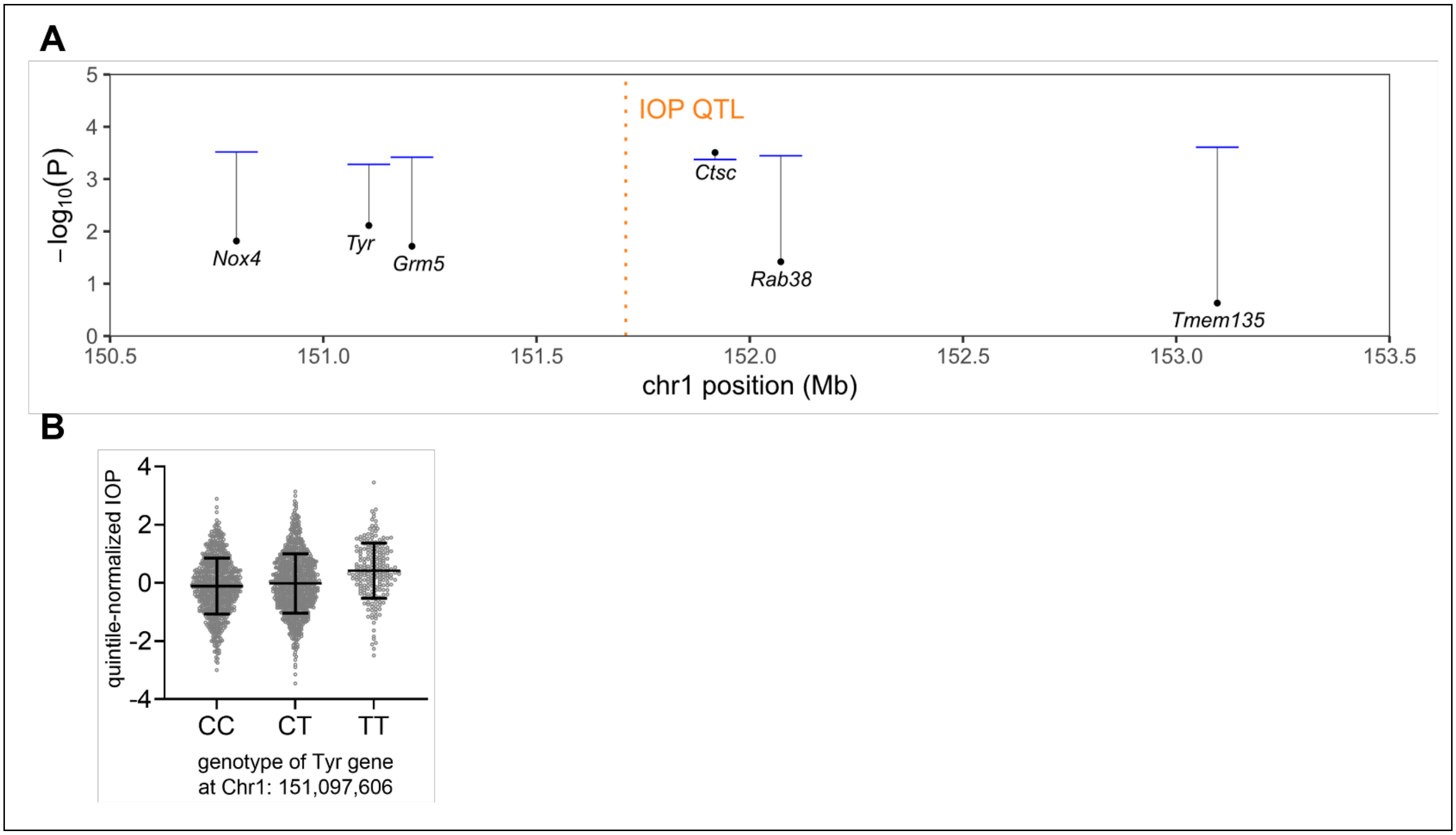


**Supplemental Figure 2.** Additional information on the QTL on Chromosome 1. (**A**) Association between gene expression and genotypes in the QTL on chromosome 1. The top-associated cis-window (+/- 0.5 Mb) SNP for expression levels of each gene within this range is indicated as a point. Each gene-specific significance threshold, determined by permutations of the gene's cis-window data, is shown as a horizontal segment connected to its top-SNP point so that a point lying below its segment indicates no significant cis-eQTL was found for that gene from whole eye tissue. (**B**) The effect plot for the *Tyr* genotype at chr1:151,097,606 shows IOP in rats with three genotypes where variant TT leads to dysfunctional protein tyrosinase and albinism in rats. The IOP is not statistically significantly different for 3 genotypes (One-way ANOVA followed by Brown-Forsythe posthoc test, p = 0.0825). IOP is shown as residuals after normalization performed as described in the "Methods" section. The number of individuals with each genotype: 726 (TT), 855 (CT), and 213 (TT).

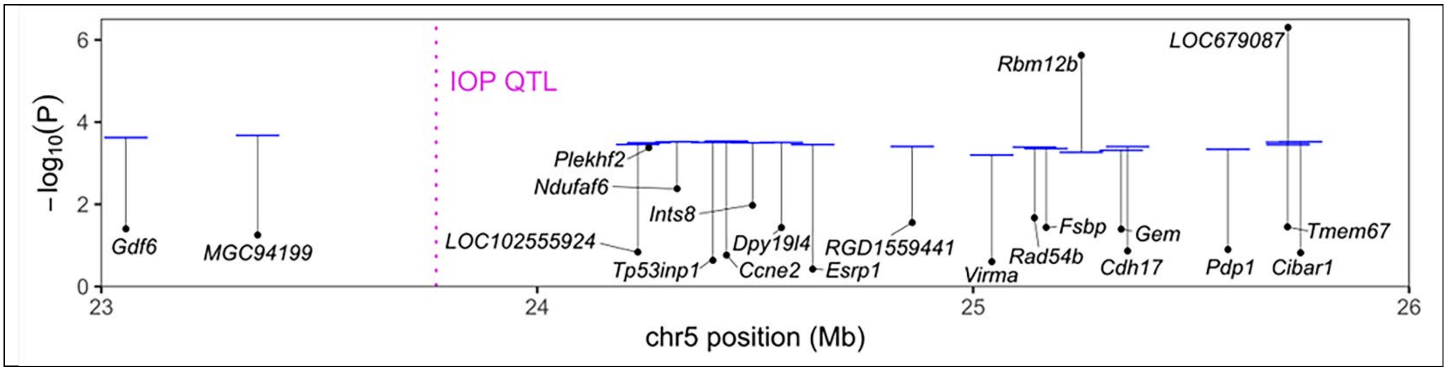


**Supplemental Figure 3.** Additional information on QTL on Chromosome 5. Association between gene expression and genotypes in the QTL on chromosome 1. The top-associated cis-window (+/- 0.5 Mb) SNP for expression levels of each gene within the 23-26 Mb range is shown as a point. Each gene-specific significance threshold, determined by permutations of the cis-window data of each gene, is shown as a horizontal segment connected to its top-SNP point, so that a point lying below its segment indicates no significant cis-eQTL was found for that gene from whole eye tissue. While both *Rbm12b* and L*OC679087* were cis-eQTLs, they are located outside of the IOP QTL region and not in LD with the QTL's top SNP. r2 between the top SNP of IOP (chr5:23768259) with the top SNPs of *Rbm12b* and *LOC679087* top (chr5:25600721 and chr5:25437533) are 0.397 and 0.395, respectively. Association analysis for *Plekhf2* expression shows that nominal *p*-value for cis-eQTL is close to the threshold.

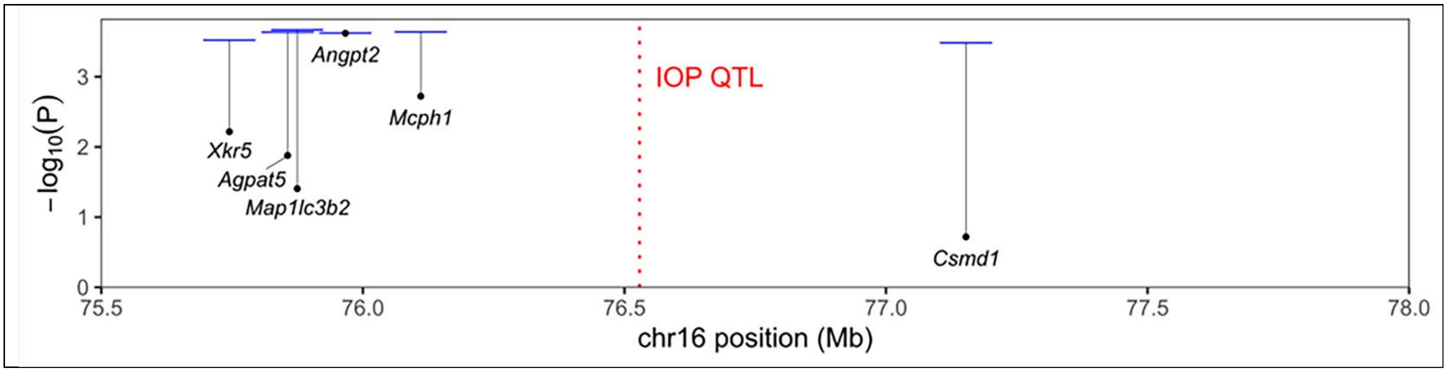


**Supplemental Figure 4.** Association between gene expression and genotypes in the QTL on chromosome 16. The top-associated cis-window (+/- 0.5 Mb) SNP for expression levels of each gene within the QTL range is shown as a point. Each gene-specific significance threshold, determined by permutations of the gene's cis-window data, is shown as a horizontal segment connected to its top-SNP point, so that a point lying below its segment indicates no significant cis-eQTL was found for that gene from whole eye tissue. The p-value for the *Angpt2* eQTL is at a threshold for significance.

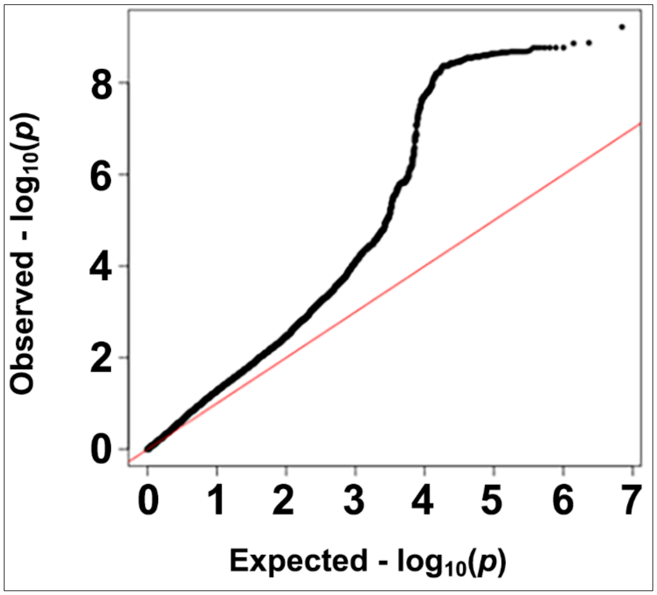


**Supplemental Figure 5.** The QQ plot for IOP GWAS.
